# Genome-wide RNAi screen for context-dependent tumor suppressors identified using *in vivo* models for neoplasia in *Drosophila*

**DOI:** 10.1101/643221

**Authors:** Casper Groth, Pooja Vaid, Aditi Khatpe, Nelchi Prashali, Avantika Ahiya, Diana Andrejeva, Madhumita Chakladar, Sanket Nagarkar, Rachel Paul, Teresa Eichenlaub, Hector Herranz, TS Sridhar, Stephen M. Cohen, LS Shashidhara

## Abstract

Genetic approaches in *Drosophila* have successfully identified many genes involved in regulation of growth control as well as genetic interactions relevant to the initiation and progression of cancer *in vivo*. Here, we report on large-scale RNAi-based screens to identify potential tumor suppressor genes that interact with known cancer-drivers: The Epidermal Growth Factor Receptor and the Hippo pathway transcriptional cofactor Yorkie. These screens were designed to identify genes whose depletion drove tissue expressing EGFR or Yki from a state of benign overgrowth into neoplastic transformation *in vivo*. We also report on an independent screen aimed to identify genes whose depletion suppressed formation of neoplastic tumors in an existing EGFR-dependent neoplasia model. Many of the positives identified here are known to be functional in growth control pathways. We also find a number of novel connections to Yki and EGFR driven tissue growth, mostly unique to one of the two. Thus, resources provided here would be useful to all researchers who study negative regulators of growth in the context of activated EGFR and/or Yki and positive regulators of growth in the context of activated EGFR.

## Introduction

Studies in genetic models of tissue growth have identified networks of signalling pathways that cooperate to control growth during animal development (reviewed in [1, 2]). Normal tissue growth involves controlling the rates of cell proliferation and cell death, as well as cell size, cell shape, etc. Signalling pathways mediate hormonal and neuroendocrine regulation of growth, which depend on nutritional status. Cell interactions also contribute to coordinating growth of cells within a tissue.

Growth regulatory pathways include both positive and negative elements to allow for feedback regulation. These feedback systems confer robustness to deal with intrinsic biological noise, and with a fluctuating external environment [3]. They also provide the means for different regulatory pathways to interact [4, 5] [6]. In the context of tumor formation, this robustness is reflected in the difficulty in generating significant mis-regulation of growth - a two-fold change in expression of many growth regulators seldom has a substantial effect on tissue size in *Drosophila* genetic models. More striking is the difficulty in transitioning from benign overgrowth to neoplasia: hyperplasia does not normally lead to neoplasia without additional genetic alterations (eg [7–9]).

Cancers typically show mis-regulation of multiple growth regulatory pathways. Mutational changes and changes in gene expression status contribute to driving cell proliferation, overcoming cell death and cellular senescence, as well as to allowing cells to evade the checkpoints that normally serve to eliminate aberrant cells. These changes alter the normal balance of cellular regulatory mechanisms, from initial cellular transformation through disease progression [10, 11]. For many tumor types, specific mutations have been identified as potent cancer drivers, with well-defined roles in disease [12, 13]. However, most human tumors carry hundreds of mutations, whose functional relevance is unknown. The spectrum of mutation varies from patient to patient, and also within different parts of the same tumor [14]. Evidence is emerging that some of these genetic variants can cooperate with known cancer drivers during cellular transformation or disease progression. The mutational landscape of an individual tumor is likely to contain conditional oncogenes or tumor suppressors that modulate important cellular regulatory networks.

Sequence-based approaches used to identify cancer genes favor those with large individual effects that stand out from the ‘background noise’ of the mutational landscape in individual cancers [10, 11]. *In vivo* experimental approaches are needed to assign function to candidate cancer genes identified by tumor genome sequencing, and to identify functionally significant contributions of genes that have not attracted notice in genomics studies due to low mutational frequency, or due to changes in activity not associated with mutation. *In vivo* functional screens using transposon mutagenesis of the mouse genome have begun to identify mutations that cooperate with known cancer driver mutations, such as K-Ras, in specific tumor models [15–17]. Genetic approaches using *Drosophila* models of oncogene cooperation have also been used to identify genes that act together with known cancer drivers in tumor formation [8, 9, 18–20] [21, 22] [2, 23]. The simplicity of the *Drosophila* genome, coupled with the ease of large-scale genetic screens and the high degree of conservation of major signaling pathways with humans, make *Drosophila* an interesting model to identify novel cancer genes and to study the cellular and molecular mechanisms that underlie tumor formation *in vivo* (reviewed in [24–27]).

In *Drosophila*, overexpression of the Epidermal Growth Factor Receptor, EGFR, or Yorkie (Yki, the fly ortholog of the YAP oncoprotein) cause benign tissue over-growth [4, 7, 9]. Combining these with additional genetic alterations can lead to neoplastic transformation and eventually metastasis [8, 9, 21, 22] [28]. Here, we report results of large-scale screens combining UAS-RNAi transgenes with EGFR or Yki expression to identify negative regulators of these growth regulatory networks that can lead to aggressive tumor formation *in vivo*. We also performed an independent screen to identify factors that could suppress EGFR-driven neoplasia. These screens have identified an expanded genomic repertoire of potential tumor suppressors that cooperate with EGFR or Yki. Interestingly, there was limited overlap among the genes that cooperated with EGFR and those that cooperated with Yki.

## Results

Overexpression of EGFR or Yki proteins in the *Drosophila* wing imaginal disc produces tissue overgrowth. Under these conditions the imaginal discs retain normal epithelial organization, but grow considerably larger than normal. However, in combination with additional genetic or environmental changes, the tissue can become neoplastic and form malignant tumors [8, 9, 22] [28]. In this context, we carried out large-scale screens using UAS-RNAi lines to identify genes which would drive hyperplastic growth to neoplastic transformation when down-regulated. To facilitate screening for tumorous growth, we expressed UAS-GFP with UAS-EGFR or UAS-Yki to allow imaginal disc size to be scored in the intact 3^rd^ instar larva (Figure 1A; screen design, examples and quality controls are shown in Supplemental Figure S1).

**Figure 1:**
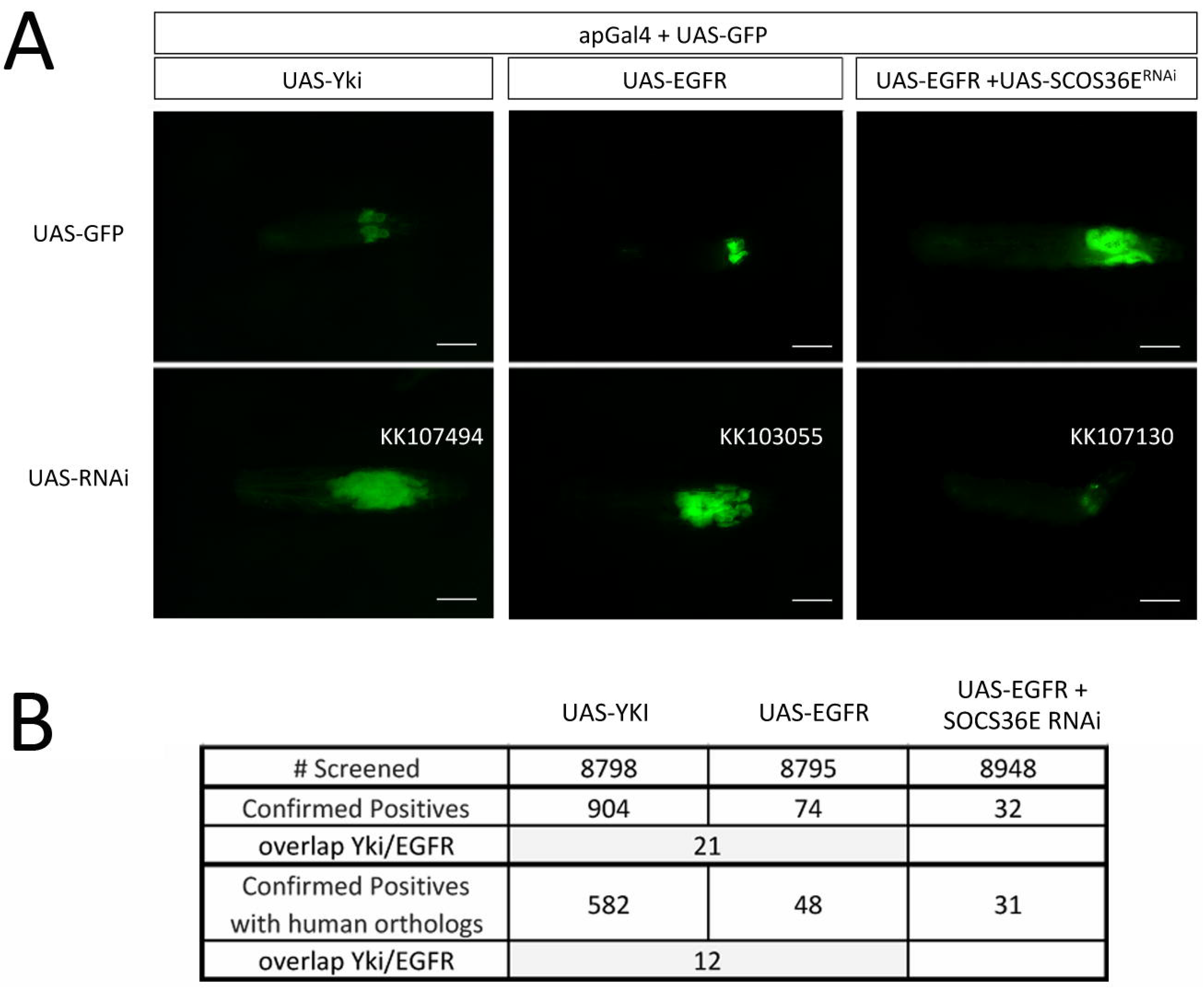
tumor formation/suppression visualized in intact larvae (A) Larvae co-expressed UAS-GFP with the indicated transgenes to permit visualization of the imaginal discs in the intact animal. All samples carried the *ap*-Gal4 driver and UAS-GFP. In addition, they carried either a second copy of UAS-GFP or one of the following: UAS-Yki, UAS-EGFR or UAS-EGFR+UAS-SOCS36E-RNAi (B) Table summarizing the number of interacting RNAi lines identified in the three large-scale screens.

A large panel of independent UAS-RNAi lines were tested for their effects on tissue growth in the EGFR and Yki expression backgrounds (Figure 1B). Hereafer, the two screens are termed as EGFR screen and Yki screen. Of ~8800 lines tested, 74 interacted with EGFR to produce tumors (~1%), whereas 904 interacted with Yki (~10%). There was limited overlap, with only 21 RNAi lines producing tumors in both screens (Figure 1B). In a parallel screen, we started with neoplastic tumors produced by co-expression of UAS-EGFR and UAS-SOCS36E-RNAi [8] and asked whether including expression of another RNAi transgene could suppress neoplasia (Figure 1A, right panels). Hereafer, this screene is termed SOCS screen. Of ~8900 lines tested, 32 suppressed tumor formation in this assay (Figure 1B). Supplemental Table S1(A) lists the genes identified in these three screens. In previous studies, massive disc overgrowth as in Figure 1(A) was often associated with loss of apically-localized Actin and E-Cadherin: features indicative of Epithelial Mesenchymal Transition (EMT); and with formation of malignant transplantable tumors [8, 9] [22]. Apico-basal polarity and Matrix Metalloprotease 1 (MMP1) expression were assessed for a randomly selected subset of lines from the EGFR and Yki screens to assess neoplastic transformation (Figure S1 and Table S1 – second sheet).

To identify the processes and pathways responsible for the interaction with the screen drivers, we looked for over-representation of biological functions among the screen positives using gene set enrichment analysis and the KEGG, REACTOME, GO and PANTHER databases. Figure 2 presents the results of the enrichment analysis as graphical interaction maps, with similar biological processes color-coded. Edge length represents similarity between genes associated with significantly enriched terms. Thus, similar terms are closer together and form a community of biological process. The genes in each cluster are shown in Figure 2 and listed in Supplemental Table S2 (has three sheets, once each for EGFR, SOCS and Yki screens).

**Figure 2:**
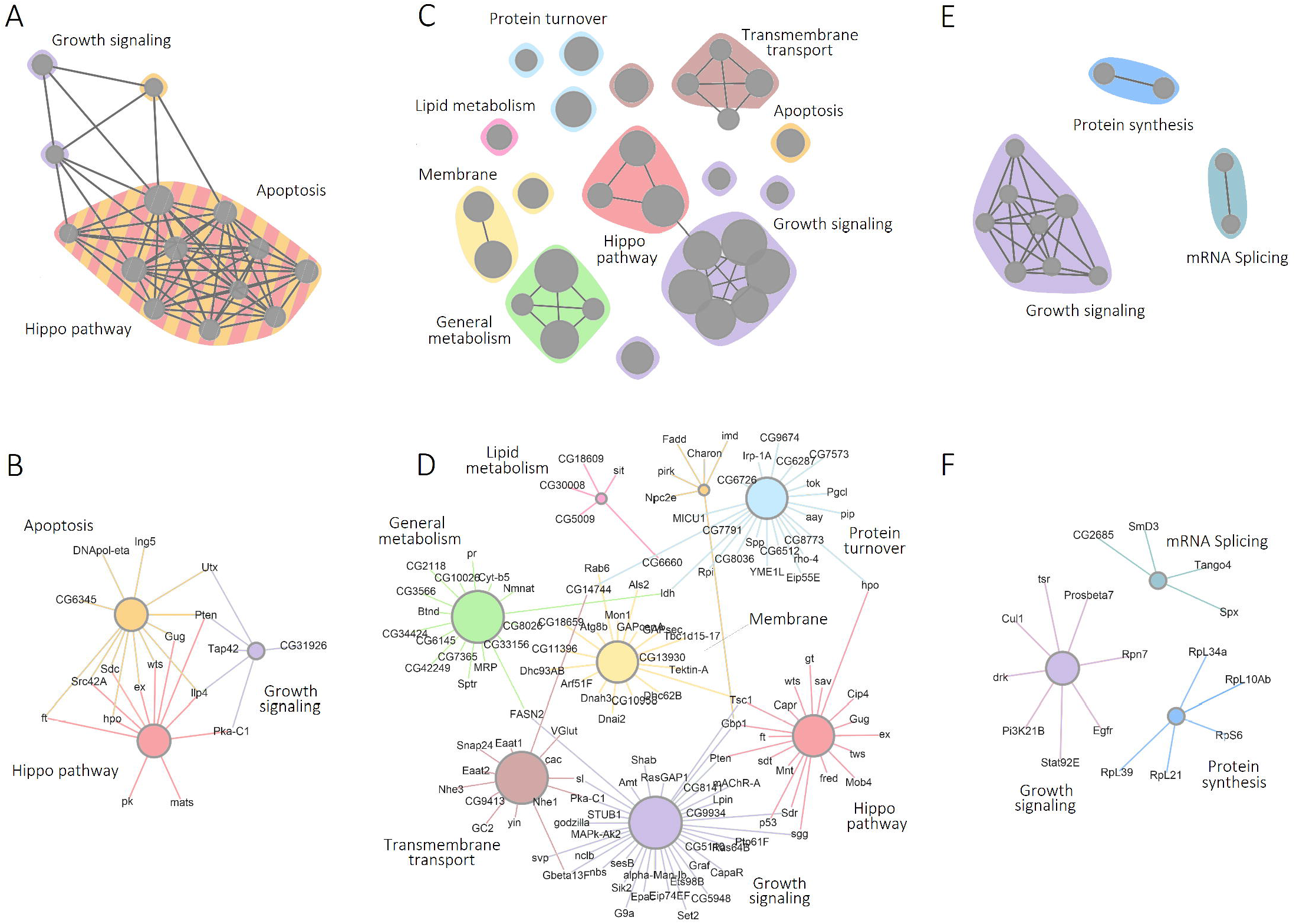
Summary of pathway enrichment analysis of fly genes identify in the *in vivo* screens reported here. (A, C, E) The results of the pathway and gene set enrichment analysis are shown as graphical interaction maps. Each node represents a significantly enriched term or pathway from the GO, KEGG, Reactome and Panther databases (Table S2). Color-coding indicates functionally related groups of terms. Lines indicate genes shared among different terms. (B, D, F) show the individual genes associated with functionally enriched cluster. (A, B) UAS-EGFR screen (C, D) UAS-Yki screen (E, F) UAS-EGFR+UAS-SOCS36E-RNAi screen

### Genes that potentially modulate EGFR function during growth control

For discs overexpressing EGFR, we observed enrichment of RNAi lines targeting the Hippo pathway, growth signaling, and apoptosis (Figure 2A, B). Many of the genes in the Hippo pathway act as negative regulators of tissue growth, so their depletion by RNAi is expected to promote growth. The Hippo pathway is known to interact with the EGFR pathway to regulate normal developmental growth [4, 5] [6]. The Hippo pathway hits included core elements of the pathway, *hpo*, *wts* and *mats*, which serve as negative growth regulators; the upstream pathway regulators *fat* (an atypical cadherin) and *expanded*; as well as the transcriptional corepressor *grunge*, which is linked to Hippo pathway activity (Table S2). Several of these loci also contributed to the enrichment of terms linked to apoptosis, along with *pten*, a phospholipase that serves as a negative regulator of PI3K/AKT signaling, *protein kinase A-C1, Src42A*, the insulin-like peptide, *ilp4*, which are also linked to growth control (Table S2).

For suppression of tumors in discs overexpressing EGFR together with SOCS36E RNAi, we observed enrichment of RNAi lines targeting signaling pathways related to growth, including elements of the AKT/PI3K pathway (Figure 2E, F, Table S2). These pathways may be required for neoplasia in this EGFR driven tumor model. As would be expected, depletion of *Egfr* limited tumor growth. Also enriched was a set of genes involved in protein synthesis (Table S2). This may reflect a need for active cellular growth machinery to support tumor growth. The significance of genes involved in RNA splicing merits further investigation.

### Genes that potentially modulate Yki function during growth control

For discs overexpressing Yki, RNAi lines targeting the Hippo pathway and associated growth regulators led to tumor production (Figure 2C, D, Table S2). These include *hpo*, *sav*, *wts, mats, ft, ex* and *gug*. Although this has not been observed previously in *Drosophila*, it is worth noting that overexpression of YAP has been shown to lead to neoplasia in mouse liver and intestinal epithelial models [29, 30]. While most cancers appear to result from activation/inactivation of multiple genes and pathways, sufficient activation of the Hippo pathway can result in neoplasia (Yki in *Drosophila* and YAP in mouse).

The Hippo tumor suppressor pathway is regulated by cell polarity, cell contact, and mechanical forces [31–33] as well as by other growth signaling pathways. Growth signaling pathways involving the *sgg*, *pten, PKA-C1, TSC1* genes among others, were also identified. Additionally, a number of genes linked to membrane-cytoskeleton interaction and transmembrane transport were found to interact, including Arf and Rab family members. We also noted the enrichment of terms related to lipid and general metabolism. Regulation of lipid metabolism might affect the properties of cellular membranes. An intriguing subgroup contain genes related to glutamatergic signaling, including the vesicular glutamate transporter *VGlut* and the *Eaat* plasma membrane glutamate transporters. The significance of these is unclear.

The large number of Yki interactors could reflect greater sensitivity of the screen. Alternatively, it might indicate a high false positive rate. While this screen was in progress, Vissers et al. [34], reported that some of the RNAi lines from the Vienna *Drosophila* RNAi KK library have the potential to produce false positives in screens based on sensitized Hippo pathway phenotypes. This proved to be due to the presence of a second transgene landing site at 40D that was found in a subset of KK lines, in addition to the 30B landing site [34, 35]. We tested the 40D landing site strain [34] and found that it did not cause a tumor phenotype under the conditions used for the screen. Nonetheless, we sampled the 40D status for a large subset of our Yki interactors (Table S1, 734/904) and found that 45% of them had insertions at 40D. A small survey comparing KK lines with Trip and GD lines showed that 65% of genes for which the KK line had a 40D site retested positive for interaction with Yki using an independent (non-KK) transgene (15/23). The Yki-interaction screen should therefore be viewed as a more sensitized sampling of potential interactors, compared to the EGFR-interaction screen.

### STRING Interactome analyses

To view all genes identified in the three screens as one functional unit (for the fact that they were all growth regulators in one or the other contexts), we made use of STRING v10 [36] to produce protein interaction maps. STRING v10 builds interaction maps by combining experimental data (including protein interaction data) with information about functional associations from text mining. STRING v10 also uses information of co-occurrence, co-expression, gene neighborhood, gene fusion, and does sequence similarity search to predict functional interaction between proteins. An interaction pair supported by multiple lines of evidence has higher confidence score than other pairs.

Figure 3A shows the STRING interaction map for the genes identified as interactors of EGFR. As noted above, Hippo pathway (red) components were prominent among the genes identified as cooperating with EGFR to drive tumor formation. Figure 3(B) shows the interaction map for the genes identified as interactors of Yki. The larger number of hits in this screen results in a more complex interaction map, with multiple interconnected clusters. The Hippo pathway (red) was again prominent in the fly screen. We also noted clusters containing elements of the ubiquitin mediated proteolysis pathway (green) and the PI3K/TOR (blue). As noted above, the higher sensitivity of this screen leads to the inclusion of weaker interactors, which may add to the complexity of these interaction maps. A focus on the stronger clusters and the interaction between them should guide future studies. Fig. 3(C) shows interaction map for the genes identified as interactors of EGFR in the suppressor screen (in discs overexpressing EGFR together with SOCS36E RNAi). Among fly genes, as expected, we observed suppression of the tumor phenotype when components of EGFR pathway are down regulated.

**Figure 3:**
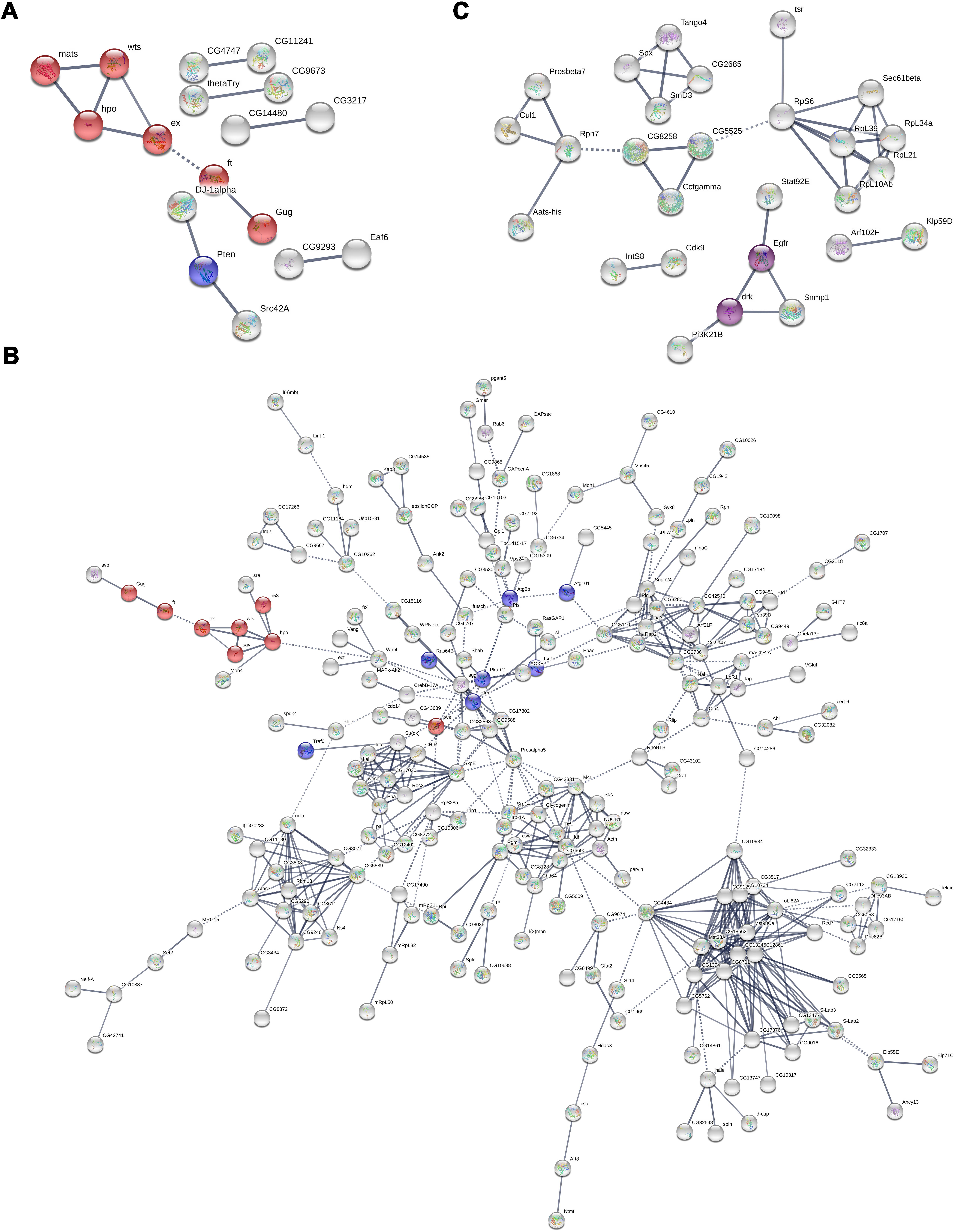
STRING interactome analysis of potential interactors of EGFR and YKi in *Drosophila*. STRING analysis was performed with confidence score of 0.7 and MCL clustering value of 2. (A) STRING Interactome of 73 fly genes identified as potential negative regulators in the context of over expression of EGFR. 17 out of those formed molecular clusters (with PPI enrichment value of 0.000482), largest being a cluster of 6 genes, all of which are constitutes of Fat/Hippo pathway (shown in red; FDR-1.39E^−5^). (B) STRING Interactome of 888 genes of identified as potential negative regulators in the context of over expression of Yki. 228 of those formed a single cluster with PPI enrichment value 1.4E-06. Components of Fat/Hippo pathway (red: FDR-0.00076) and Autophagy genes (blue: FDR-0.0241) are enriched in this cluster. (C) STRING Interactome of 32 fly genes identified as potential oncogenes in the context of SOCS suppression. 27 out of those formed molecular clusters (with PPI enrichment value of 0.0122), largest being a cluster of 14 genes. A smaller cluster comprising of EGFR and DrK were enriched in Dorso-ventral axis formation (shown in purple: FDR-0.0089).

### Human orthologs of the fly genes identified in the three screens

To identify human orthologs for the candidate genes, we used the DRSC Integrative Ortholog Prediction Tool, DIOPT (Version 7.1, March 2018; www.flybase.org). DIOPT scores reflect the number of independent prediction tools that identify an ortholog for a given *Drosophila* gene. Orthology relationships are usually unambiguous when found by most of the 12 independent prediction tools in DIOPT. Table S1 lists the primary human orthologs (highest weighted DIOPT score), as well as the other orthologs with a weighted DIOPT score >2 for each of the hits in the fly screen. The primary human ortholog was used for subsequent analysis. In cases where multiple human orthologs had the same score, all orthologs with highest weighted DIOPT score were used. Out of 73 EGFR positive hits, 46 genes had one or more human orthologs, in total mapping to 50 human genes. Out of 32 SOCS positive hits 30 genes had one or more human orthologs, in total mapping to 31 human genes. Out of 904 YAP positive hits 570 genes had one or more human orthologs, in total mapping to 611 human genes.

To view the human orthologs in a functional context, we performed a gene set enrichment analysis and the KEGG, REACTOME, GO, PANTHER, NCI, MsigDB, BIOCARTA databases. Figure 4 presents the results of the enrichment analyses as graphical interaction maps, with similar biological processes color-coded. Edge length represents similarity between genes associated with significantly enriched terms. Thus, similar terms are closer together and form a community of biological processes. The genes in each cluster are shown in Figure 4 and listed in Supplemental Table S3. Because the enrichment analysis is highly sensitive to the number of orthologs for each of the fly genes, we used the minimal set consisting of only the primary human orthologs.

**Figure 4:**
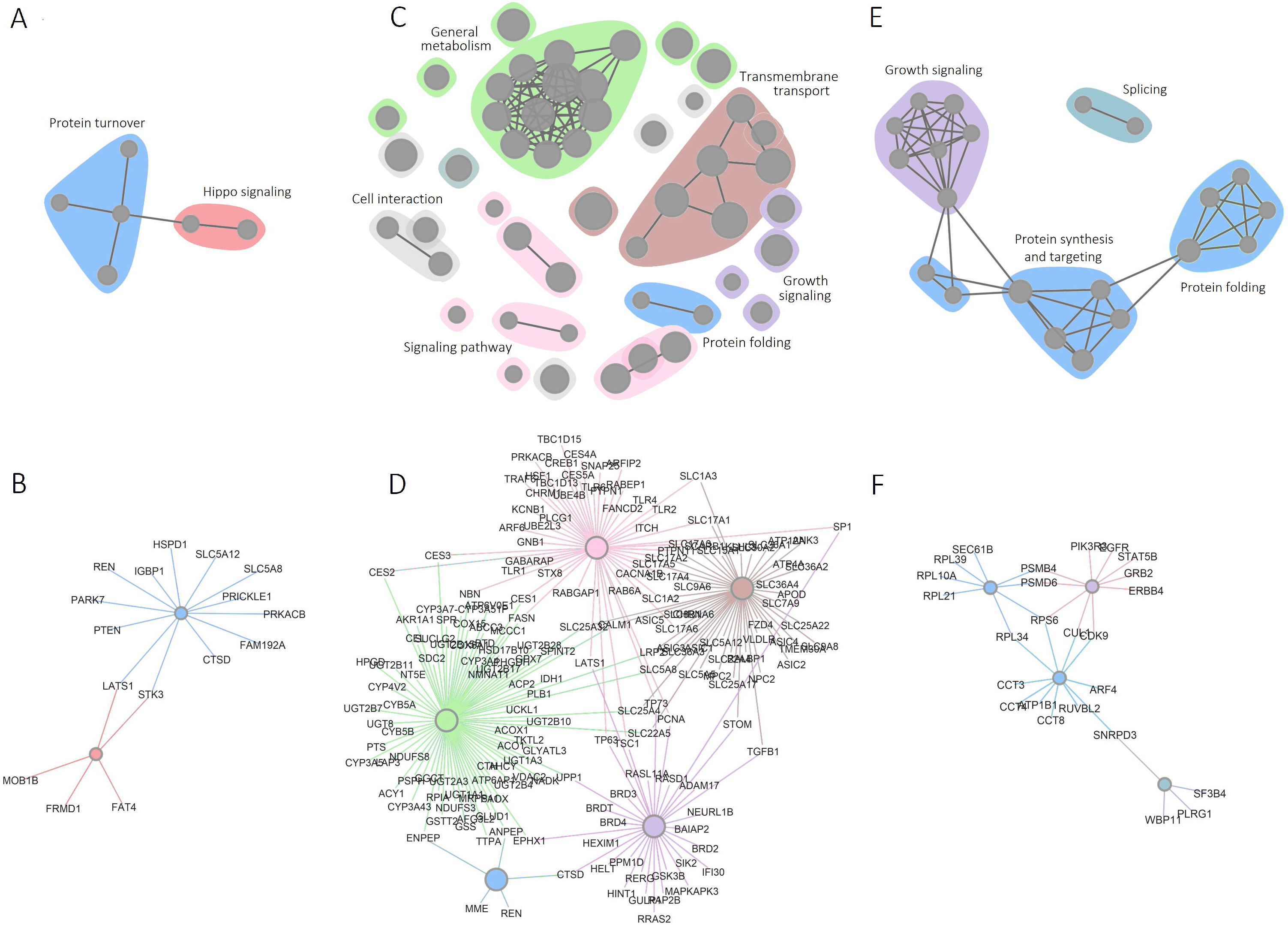
Summary of pathway enrichment analysis of human orthologs (A, C, E) The results of the pathway and gene set enrichment analysis are shown as enrichment maps. Each node represents a significantly enriched term or pathway from the GO, KEGG, Reactome and PANTHER, NCI, MsigDB, BIOCARTA databases (Table S3). Color-coding indicates functionally related groups of terms. Lines indicate genes shared among different terms. (B, D, F) show the individual genes associated with functionally enriched cluster. (A, B) UAS-EGFR screen (C, D) UAS-Yki screen (E, F) UAS-EGFR+UAS-SOCS36E-RNAi screen

Hippo pathway components were enriched among the orthologs cooperating with EGFR to drive tumor formation (Fig 4A, B; Table S3). Two of these, LATS1 and STK3, also contributed to enrichment for a term linked to protein turnover. Regulation of protein turnover is an important mechanism for controlling the activity of a number of Hippo pathway components. For the screen for suppression of tumors in discs overexpressing EGFR together with SOCS36E RNAi, we observed enrichment of orthologs targeting growth signaling pathways, protein synthesis and mRNA splicing (Figure 4E, F, Table S3), similar to what was seen for the fly gene set analysis. We also observed enrichment of pathways related to protein folding and molecular chaperones, in the human gene set. For the Yki screen, the human ortholog set was enriched for terms related to general metabolism, and membrane transport, as well as growth signaling, and other signaling pathways, including genes involved in protein turnover (Fig 4C, D).

## Discussion

The Hippo pathway has emerged from this study as the single most important pathway limiting tumor formation in *Drosophila*. Increasing Yki activity by depletion of upstream negative regulators promoted tumor formation in both the EGFR and Yki hyperplasia models. Yki controls tissue growth by promoting cell proliferation and by concurrently inhibiting cell death through targets including *Diap1*, *CycE* and *bantam* miRNA [7, 36–39]. The central role of the Hippo pathway as an integrator of other growth-related signals may also contribute to the abundance of tumor suppressors associated with Yki-driven growth [1, 2, 26]. Mis-regulation of this pathway also contributes to tumor formation in mouse models [40].

The potential of Yki/YAP expression to drive cellular transformation has been highlighted by studies of primary human cells in culture, which have shown that YAP expression is both necessary and sufficient to confer a transformed phenotype involving anchorage independent growth and the ability to form tumors in xenograft models [41, 42]. We therefore consider it likely that the consequence of Yki overexpression predispose the tissue to transformation, allowing identification of a richer repertoire of cooperating factors. Indeed, YAP overexpression has been causally linked to formation of specific human tumors [43, 44]. The Hippo pathway has also been implicated in tumor formation resulting from cytokinesis failure [45] and this has recently been linked to Yki-mediated regulation of *string (*CDC25) expression [46]. The sensitivity of Yki-expressing tissue to tumor formation might be explained by the finding that Yki promotes cell cycle progression at both the G1-S transition (through regulation of *cycE* [7] and at the G2-M transition through regulation of *string*. In contrast, mitogens and growth factors such as EGFR typically induce growth by promoting G1-S, and therefore remain somewhat constrained by the G2-M checkpoint.

While our manuscript was in preparation, another group reported an RNAi screen to identify loci cooperating in tumorigenesis driven by expression in eye discs of the oncogenic activated mutant form of *Ras* [47]. We note that the activated Ras RNAi screen produced over 900 hits, compared with 74 for our EGFR screen, suggesting that the Ras screen was considerably more sensitized. We were surprised to note that there was almost no overlap between the two screens. This suggests that the genetic interactions required to promote tumorigenesis in the context of expression of an activated mutant form of RAS are distinct from those required to promote tumorigenesis in the context of native EGRF overexpression. And perhaps, the differences between the tissue contexts (eye discs in [47] vs wing discs in our screen). It will be of interest, in future, to learn whether this distinction holds true for factors promoting tumor formation in human cancers that depend on EGFR overexpression vs those dependent on Ras mutants.

To conclude, the results reported here provide an extensive assessment of the genes that can serve as negative regulators of growth that can contribute to the formation of neoplastic tumors *in vivo* in *Drosophila*. In addition to finding genes linked to known growth control pathways, a number of novel connections to Yki and EGFR driven tissue growth have been identified, which merit further investigation in the *Drosophila* genetic model. Exploring the potential relevance of genes identified in this manner to human cancer will involve assessing the correlation of candidate gene expression with clinical outcome across a broad range of cancers (eg [28, 48]), as a starting point to identify biomarkers as well as novel candidate drug targets.

## Materials and Methods

### RNAi Screens

The KK transgenic RNAi stock library was obtained from the Vienna *Drosophila* RNAi Center (www.vdrc.at) carrying inducible UAS-RNAi constructs on Chromosome II. For each cross, 5 males from the KK transgenic RNAi stock were crossed separately to 10-15 virgins from each of the following three driver stocks: *w*, ap*-Gal4, UAS-*GFP*/CyO; UAS-*Yki*, *tub*-Gal80^**ts**^/TM6B (Yki driver); *w\**; *ap*-Gal4, UAS-*GFP*/CyO; UAS-*EGFR*, *tub*-Gal80^**ts**^/TM6B (EGFR driver); and *w\**; *ap*-Gal4, UAS-*GFP*/CyO; and *w\**; *ap*-Gal4, UAS-*GFP*, *Socs36E*^**RNAi**^/CyO; UAS-*EGFR*, *tub*-Gal80^ts^/TM6B (EGFR driver +SOCS36E RNAi).

Virgin female flies were collected over 4-5 days and stored at 18°C in temperature-controlled incubators on medium supplemented with dry yeast, prior to setting up crosses. Virgin females were mated to KK stock males (day 1) and the crosses were stored at 18°C for 4 days to provide ample time for mating before starting the timed rearing protocol used for the screen. On day 5, the crosses were transferred into new, freshly yeasted vials for another 3 days at 18°C. On day 8, the adult flies were discarded, and larvae were allowed to develop until day 11, at which time the vials were moved to 29°C incubators to induce Gal4 driver activity. Crosses were aged at 29°C for a further 8-9 days, after which larvae were scored for size and wing disc overgrowth phenotypes for Yki and EGFR driver screen crosses. Flies were scored for suppression of the tumor phenotype for the EGFR driver +SOCS36E RNAi crosses (see Supplemental Fig. S1 for the screen design and workflow).

In order to verify the integrity of the driver stocks during the course of the screen, we examined their expression patterns in conjunction with setting up screen crosses each week. For each driver, 2-3 of the bottles used for virgin collection were induced at 29°C for 24 hours and analyzed using fluorescence microscopy for *apterous-*Gal4 specific expression in wandering 3-instar larvae.

Positive hits form the initial screen were retested by setting up 2 or more additional crosses. The hits were scored as verified if 2 out of 3 tests scored positive. Wandering third instar larvae of confirmed positives were imaged and documented using fluorescence microscopy.

### Genomic DNA PCR 40D landing site occupancy test

Genomic DNA from a select number of *Drosophila* KK transgenic RNAi library stocks was isolated following a protocol available at the VDRC (www.vdrc.at). The presence or absence of the KK RNAi transgene at the 40D insertion site on the second chromosome was determined by multiplex PCR using the following primers:

40D primer (C_Genomic_F): 5’-GCCCACTGTCAGCTCTCAAC-3’ pKC26_R: 5’-TGTAAAACGACGGCCAGT-3’

pKC43_R: 5’-TCGCTCGTTGCAGAATAGTCC-3’

PCR amplification was performed using GoTaq G2 Hot Start Green Master Mix kit (Promega) in a 25 µL standard reaction mix and the following program: initial denaturation at 95°C for 2 min, followed by 33 cycles with denaturation at 95°C for 15 sec, annealing at 58°C for 15 sec and extension at 72°C for 90 sec. One final extension reaction was carried out at 72°C for 10 min. Reactions were stored at −20°C prior to gel loading. PCR using these primers generate an approximately 450 bp product in case of a transgene insertion or a 1050 bp product in case of no transgene insertion site at 40D.

### Screen Database

Results from the three screening projects were added to a screen management database, www.iiserpune.ac.in/rnai/, including images of positive hits and background information such as RNAi line ID, corresponding gene information from the Flybase etc. The database was developed by Livetek Software Consultant Services (Pune, Maharashtra, INDIA).

### Pathway and gene set enrichment analysis

Gene set enrichment analysis was performed using genes that upon down regulation induced tumor formation (EGFR, YKI background) or suppressed tumor formation (EGFR+SOCS background). For *D. melanogaster* enrichment analysis all *D. melanogaster* protein coding genes were used as the “gene universe” together with organism specific datasets. For human ortholog enrichment analysis all human protein coding genes were used as the “gene universe” together with organism specific datasets. The algorithm packages and databases used in analysis are listed in Supplemental Tables S2 and S3. Unless otherwise specified, pathway databases included in these packages were used. The KEGG database was downloaded directly from source on 10.10.2018. Organ system specific and disease related pathway maps were excluded from this analysis. Minimum and maximum number of genes per pathway or gene set, significant criteria, minimum enriched gene count and annotated gene counts for each test and database are indicated in Supplemental Tables S2 and S3. GO results were filtered for level >2, to eliminate broad high-level categories and <10 to minimize duplication among subcategories. A representative term was selected in the cases were identical set of genes mapped to multiple terms within the same database. After filtering, the top 10 terms from each database were used for clustering analysis.

Pathway and gene set enrichment analysis results were visualized as enrichment map with appropriate layout based on gene overlap ration using igraph. Gene overlap ratio was set as edge width. Edges with low overlap were deleted, filtering threshold was based on a number of “terms” in the results table – from 0 to 50 by 10; increasing filtering thresholds from 0.16 to 0.26 by 0.2. Clusters were detected using “Edge betweenness community” algorithm. Similar biological processes were color-coded.

### R packages

*clusterProfiler (3.8.1)* - [49].

*ReactomePA (1.24.0)* - [50].

http://pubs.rsc.org/en/Content/ArticleLanding/2015/MB/C5MB00663E.

*graphite (1.26.1)* - Sales G, Calura E, Romualdi C (2018). graphite: GRAPH Interaction from pathway Topological Environment. R package version 1.26.1.

igraph (1.2.2) - Csardi G, Nepusz T: The igraph software package for complex network research, InterJournal, Complex Systems 1695. 2006. http://igraph.org

### Database references

KEGG – [51, 52].

REACTOME – [53]

Panther – [54]

GO – [55].

### STRING interaction maps

STRING v10 is a computational tool for protein interaction network and pathway analysis [56]), to identify significant functional clustering among the candidate genes. STRING builds interaction maps by combining experimental data (including protein interaction data) with information about functional associations from text mining. STRING interactome maps were used to search for statistically significant enrichment of KEGG pathways.

## Acknowledgements

The professionalism, tireless support and goodwill provided by Snehal Patil, Yashwant Pawar and Bhargavi Naik from the IISER Pune fly facility contributed greatly to the success of the screening projects reported here.

## Funding

This work was primarily supported by a Indo-Danish research grant from Department of Biotechnology, Govt. of India to TSS and LSS and from Innovationfund Denmark, Novo Nordisk Foundation NNF12OC0000552 and Neye Foundation to SMC. JC Bose Fellowship and grant from Department of Science & Technology, Govt of India to LSS. We thank other members of all the laboratories for critical input.

